# Sentieon DNAscope LongRead – A highly Accurate, Fast, and Efficient Pipeline for Germline Variant Calling from PacBio HiFi reads

**DOI:** 10.1101/2022.06.01.494452

**Authors:** Donald Freed, William J. Rowell, Aaron M. Wenger, Zhipan Li

## Abstract

PacBio^®^ HiFi sequencing is the first technology to offer economical and highly accurate long-read sequencing, with average read lengths greater than 10kb and average base accuracy of 99.8%^1^. Here, we present DNAscope LongRead, an accurate and efficient pipeline for germline variant calling from PacBio^®^ HiFi reads. DNAscope LongRead is a modification and extension of Sentieon’s DNAscope tool, a precisionFDA award-winning variant caller. DNAscope LongRead is computationally efficient, calling variants from 30x HiFi samples in under 4 hours on a 16-core machine (120 virtual core-hours) and highly accurate, with precision and recall on the most recent GIAB benchmark dataset exceeding 99.83% for HiFi samples sequenced at 30x coverage, and robust to changes in benchmark dataset and upstream library preparation and sequencing.

## Intro

Since its introduction nearly two decades ago, short-read sequencing has become commonplace in genomics and is frequently applied in both clinical and research settings. However, short read platforms have fundamental limitations due to the diploid and repetitive nature of most complex genomes, where haplotype and some location information are necessarily lost during the short read sequencing process. This is particularly problematic for clinical and research studies of the Human genome, where some known clinically-relevant genes and certain types of structural variations are almost completely inaccessible^2–8^. Long-read sequencing platforms solve many of these issues but initially were limited by higher read error rates.

In 2019, PacBio developed HiFi reads, an optimization of earlier circular consensus sequencing (CCS) methodologies^9^, to produce reads that are both long and highly accurate^1^. CCS uses hairpin adapter sequences to circularize DNA fragments. These molecules are then sequenced multiple times around the circle, generating a series of subreads. Subreads are then combined computationally to create a consensus read. Due to the unbiased nature of sequencing errors in the PacBio technology^10–12^, errors present in individual subreads can be identified and corrected, resulting in consensus reads that are more faithful representations of the original DNA molecule. Overall, the process produces reads that are both long, greater than 10kb, and highly accurate, with average base accuracy of 99.8%. The combination provides benefits to downstream applications where the higher read accuracy enables accurate variant calling. The long read length provides access to difficult regions of the genome, including medically-relevant genes that had been previously identified as inaccessible with short-read sequencing technologies^1,7^.

Sentieon Inc. develops fast, accurate, and efficient tools for genomic data processing, including replacements for the GATK best-practices pipelines for germline and somatic variant calling^13,14^. To further improve germline variant calling accuracy beyond the accuracy provided by the GATK’s HaplotypeCaller, Sentieon developed DNAscope as an improved-accuracy germline variant caller and used an earlier version of DNAscope to participate and win awards in the original PrecisionFDA (pFDA) Truth Challenge. DNAscope has a very similar architecture to the previously described architecture of the GATK’s HaplotypeCaller, but implements a more robust algorithm for local assembly, allowing variant calling across more complex and repetitive genomic regions. Sentieon also offers pre-trained machine learning models for variant genotyping, which can be used with DNAscope to achieve high accuracy human WGS germline variant calling.

Here we present an update and modification of Sentieon’s DNAscope to handle PacBio HiFi reads. For this pipeline, we tune variant calling parameters to better adapt DNAscope to the longer HiFi reads and incorporate new statistical and machine-learning models for variant calling. The updated and modified DNAscope is then incorporated into a multi-pass germline variant calling pipeline. Benchmarking of the new pipeline demonstrates a 20% accuracy improvement compared to the pFDA Truth Challenge V2 winning submission, while maintaining robust performance across diverse truthsets and samples.

## Results

### Development of a new pipeline for PacBio HiFi reads

Sentieon’s DNAscope germline variant caller uses a similar logical architecture to the GATK HaplotypeCaller, implementing software modules for active region detection, local assembly, and PairHMM. The major improvements in DNAscope are a more robust algorithm for local assembly and use of a machine-learned model for variant genotyping and filtration. These improvements work together to significantly improve variant calling recall and precision. Other minor improvements include updates to the active region detection logic and the addition of variant annotations that are useful for variant filtration.

We introduce additional changes to DNAscope to optimize variant calling for PacBio HiFi data. To better adapt DNAscope’s variant calling to the longer PacBio HiFi reads, we increase the size of detected active regions. Additionally, we introduce process-specific models for variant calling of haploid, diploid, and unphased regions of the genome.

PacBio HiFi reads have mean base accuracies exceeding 99.8%. The vast majority of errors are indels in homopolymer contexts, which creates a distribution of observed repeat lengths. For multiploid samples, the distribution of observed repeat lengths is an aggregate of distributions from individual haplotypes. To improve variant calling accuracy at these sites, we perform read-backed phasing of SNVs discovered during an initial pass of variant calling. Phased variants are then used to tag overlapping reads with their haplotype of origin. During a second pass of variant calling, phased variants are then used to tag overlapping reads with their haplotype of origin, effectively compartmentalizing the aggregate read distribution to enable more accurate variant calling across phased regions.

The adapted DNAscope and new functionality for read-backed phasing and read tagging are combined into an easy-to-use pipeline for germline variant calling from PacBio HiFi reads (Figure 1). The pipeline can be broadly divided into three stages: (1) repeat model calibration, variant candidate generation, and SNV calling from the unphased reads. (2) Variant phasing and phased repeat model calibration. (3) Haploid variant candidate generation and haploid variant calling across the phased genome and diploid variant calling across the unphased regions of the genome. The pipeline optionally includes additional handling of the MHC region that has been tuned empirically to further improve accuracy across the MHC. The pipeline is implemented in portable POSIX-compliant shell script with a command line interface created with argbash^15^.

**Figure 1.**
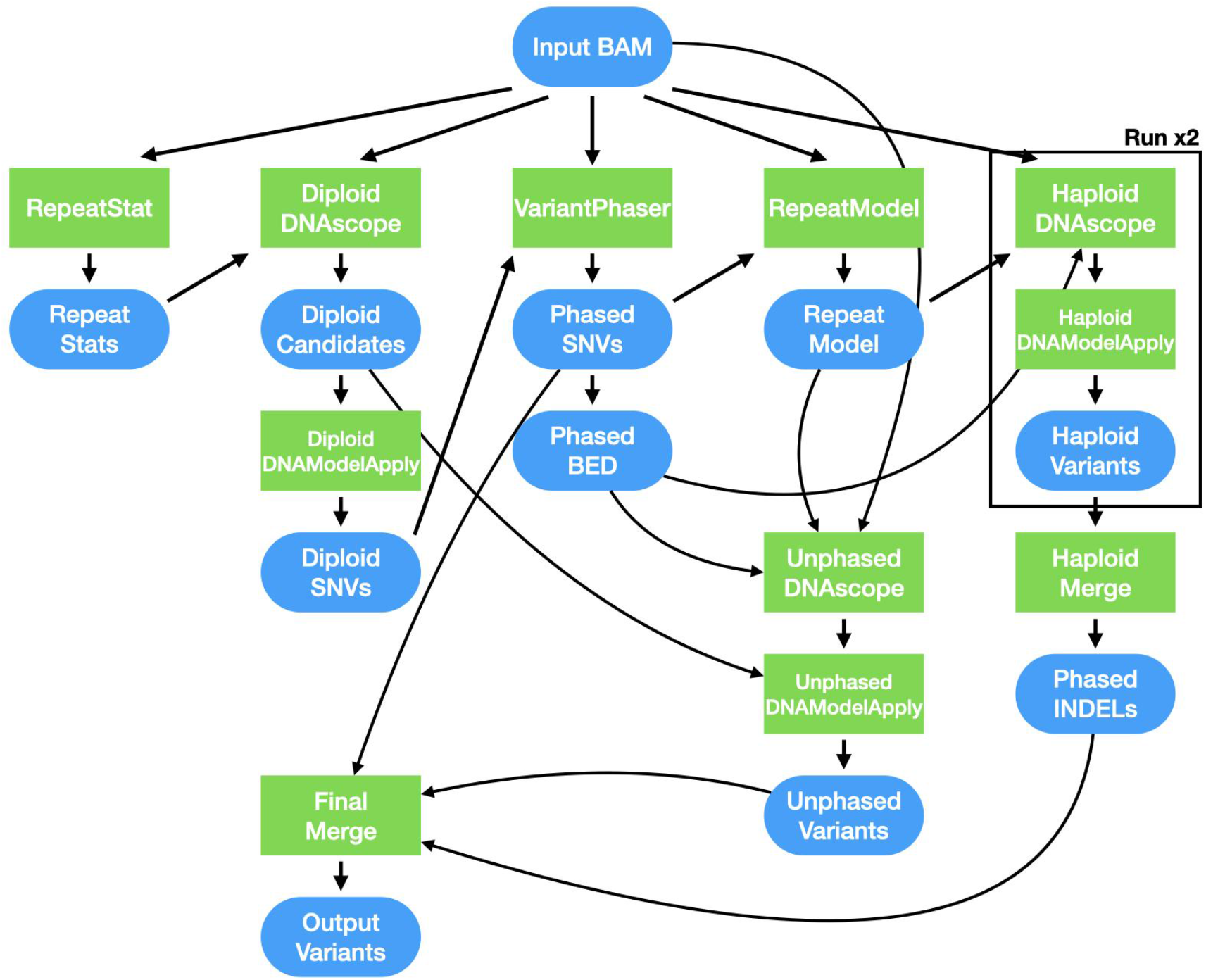
Overview of the DNAscope LongRead pipeline. The DNAscope LongRead pipeline calls germline variants from aligned PacBio HiFi reads. Conceptually, the pipeline can be divided into three phases: a first pass of variant calling, variant phasing, and a second, more accurate, pass of variant calling across phased regions of the genome, processing each haploid parental genome separately. Before the first and second passes of variant calling, statistical models are calibrated to the repeat content of the sample, improving both variant calling accuracy and pipeline robustness. The core variant calling pipeline calls DNAscope across the phased or unphased regions of the genome and uses DNAModelApply to perform model-informed variant genotyping. Small Python scripts are used for VCF manipulation.

### Accuracy on the precisionFDA Truth V2 dataset

We initially sought to evaluate the performance of our pipeline using PacBio HiFi samples sequenced for the precisionFDA (pFDA) Truth Challenge V2, the prestigious germline variant calling challenge operated by the Food and Drug Administration. pFDA challenges are open to the public and performance on the challenge provides a good measure of state-of-the-art variant calling accuracy from HiFi reads^16^.

Using the DNAscope LongRead pipeline, we called variants from the ∼35x HG002, HG003, and HG004 PacBio HiFi samples used during the challenge after alignment to GRCh38 and evaluated the variant calls using the GIAB v4.2.1 truthset. We then compared our pipeline benchmarks to the top performing pipeline for overall accuracy (Figure 2; Supplementary Table 1). DNAscope LongRead called an average of 3,859,105 variant records inside the GIAB v4.2.1 high-confidence regions, with an average of 9,130 errors across the three samples, a 15% reduction of mean errors compared to the pFDA winning pipeline for PacBio HiFi data. The mean F1-score for the three pFDA samples is 0.9988 for the DNAscope LongRead pipeline compared to 0.9986 for the pFDA winning pipeline. DNAscope LongRead performs especially well across more difficult genomic regions, including the MHC and difficult-to-map regions where mean error reductions are 55% and 27% respectively. This demonstrates how advances in variant calling can improve accuracy for a fixed set of sequencing reads.

**Figure 2.**
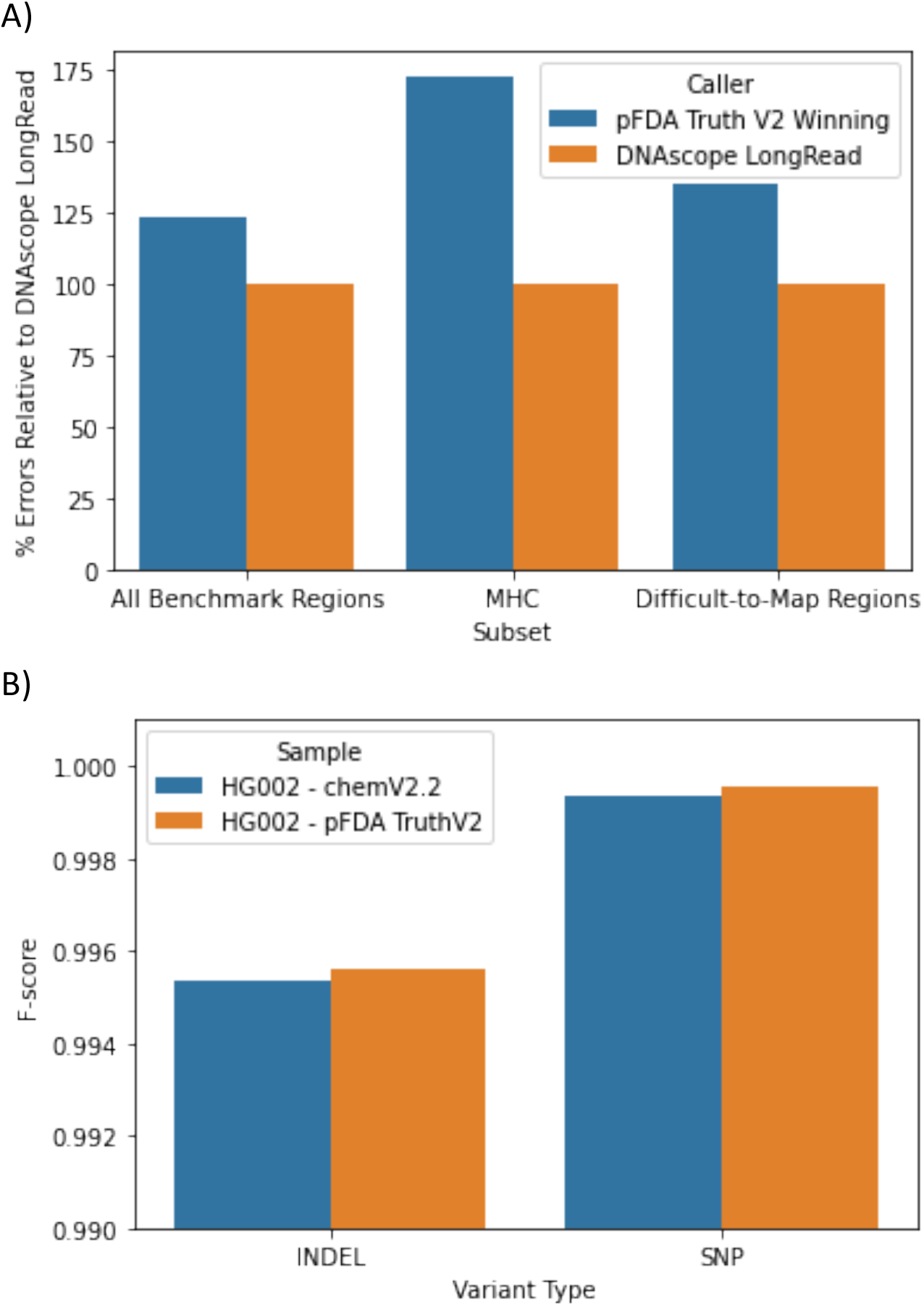
Variant calling accuracy on PacBio HiFi samples from the PrecisionFDA Truth Challenge V2 and newer chemistries. (A) The DNAscope LongRead pipeline provides a substantial reduction in the total number of errors relative to the PrecisionFDA Truth Challenge V2 winning submission. Errors are especially lower across the MHC and Difficult-to-Map stratifications. (B) The DNAscope LongRead pipeline is robust to changes in library preparation and sequencing chemistry. The chemV2.2 sample uses a new sequencing chemistry and provides comparable accuracy to the HG002 sample used in model training.

### Accuracy on new chemistries

The models incorporated into the DNAscope LongRead pipeline are trained on data from one HG001, three HG002, and two HG004 HiFi samples. Full details of samples used in model training are recorded in (Supplementary Table 3). The above experiments use samples sequenced for the pFDA Truth V2 Challenge on Sequel II system with 2.0 chemistry and consensus basecalling using SMRT Link v8.0 and ccs version 4.0.0 and demonstrate good performance with such samples^16^. To test the ability of DNAscope LongRead to extend to updated samples, we obtained samples sequenced with PacBio’s chemistry 2.2 on a Sequel II. After CCS read generation, the sample was sequenced to approximately 41x-fold coverage with HiFi reads. HiFi reads were then aligned to the Human reference genome using pbmm2, variants were called using DNAscope LongRead, and the resulting variant calls were compared to the GIAB v4.2.1 benchmark truthset for the HG002 sample.

DNAscope LongRead called a total of 9,231 errors relative to the GIAB v4.2.1 benchmark dataset with the chemistry v2.2. sample, for an overall F1-score of 0.9988, comparable to the 9,130 mean errors and 0.9988 F1-score using the pFDA Truth V2 sample chemistry v2.0 (Supplementary Figure 2, Supplementary Table 4). DNAscope extends well to newer sequencing chemistries and `ccs` versions in our testing. While we expect DNAscope LongRead’s models may need re-training as the PacBio platform continues to improve, these results help demonstrate the pipeline’s robustness to common changes in the upstream library preparation and sequencing.

### Accuracy on a serially downsampled dataset

The pFDA data represent a high-quality dataset at a good level of coverage. However, samples are often sequenced to lower coverages to allow for sequencing of additional samples. To test the robustness of our pipeline with lower coverage samples, we performed a serial downsampling of the 35x HG003 sample from the pFDA Truth V2 Challenge. The downsampling step was 5x from 5x to 35x with additional samples later added between 5x and 20x to better visualize changes in accuracy between these coverages (Figure 3, Supplementary Table 2). SNV and indel accuracy outside of long homopolymer contexts extends well to lower coverages, with a reduction in F1-score of only 0.00074 and 0.0049 between 35x and 20x coverage for SNVs and indels outside of long homopolymers, respectively. Overall indel variant calling accuracy is more affected by coverage, with a 0.013 reduction in F1-score between 35x and 20x coverage.

**Figure 3.**
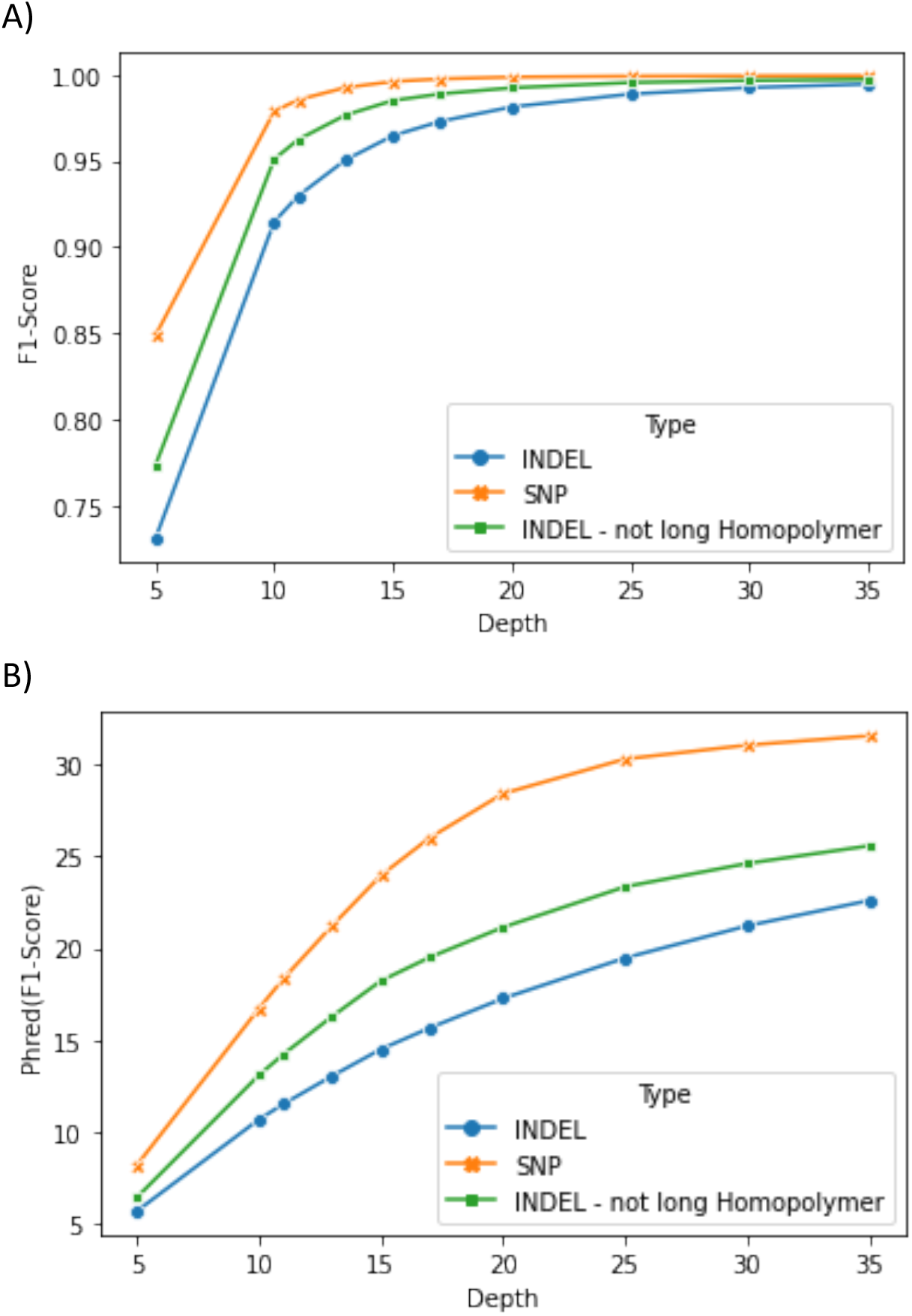
The effect of read coverage on DNAscope LongRead pipeline accuracy. F1-score for SNVs (x), indels (dots), and indels outside of long homopolymer contexts (squares) from a serial downsampling experiment. Downsampling step is 5x from 35x read coverage to 20x read coverage, while additional samples were added between 10x and 20x read coverage. SNV accuracy is more robust than indel accuracy at lower read coverages. Data is shown in two views to highlight accuracy improvements at higher read depths (A) F1-score and (B) phred-scaled F1-score.

### Accuracy on the CMRG dataset

Thus far, we have only assessed the performance of our pipeline against the GIAB v4.2.1 truthset. The Challenging Medically Relevant Genes (CMRG) benchmark was recently developed by the GIAB consortium to extend benchmark regions to medically relevant genes that are only partially covered by the GIAB v4.2.1 benchmark regions^17^. In addition to adding new genomic regions, the CMRG benchmark more heavily incorporates information from high-quality diploid assemblies. Accordingly, the CMRG benchmark provides a high-quality, but partially orthogonal benchmark relative to the GIAB v4.2.1 benchmark used for model training.

To test the ability of DNAscope LongRead to extend outside of the GIAB benchmark regions, we evaluated the variants called by DNAscope LongRead using the pFDA Truth V2 HG002 sample against the CMRG benchmark dataset (Supplementary Figure 1). Similar to the previously reported results for DeepVariant^17^, DNAscope LongRead generates 846 total errors for the sample. DNAscope LongRead correctly ignores most incorrectly mapped reads adjacent to sample-specific insertions in KMT2C. These incorrectly mapped reads are responsible for 277 false-positive variant calls in a DeepVariant-HiFi pipeline but only 62 in DNAscope, significantly reducing false-positives relative to the previously reported DeepVariant results. These results demonstrate the extensibility of the DNAscope LongRead pipeline to regions outside the GIAB v4.2.1 benchmark and to orthogonally created truthsets.

### Runtime performance and memory usage

For all invocations of the DNAscope LongRead pipeline, memory usage was profiled using Snakemake’s built-in benchmarking functionality, overall runtime was measured by the controlling shell, and per-stage runtime and memory usage metrics for most stages are reported by the Sentieon tools (Figure 4; Supplementary Tables 5 and 6). Runtime for the DNAscope LongRead pipeline ranged from 4.52 hours for the chemistry v2.2 sample of X coverage to 1.71 hours for the 5x downsample of the pFDA Truth Challenge V2 HG003 sample. Runtime for the 30x downsample of the pFDA Truth Challenge V2 HG003 sample was 3.76 hours, or 120.3 core-hours. Higher read coverages were associated with longer runtimes and runtime was relatively consistent across samples with a similar level of read coverage. Although not explicitly tested in this study, the DNAscope LongRead pipeline scales well to larger servers with higher thread counts and much faster turnaround times under 40 minutes can be achieved on larger servers with more recent CPU architectures.

**Figure 4.**
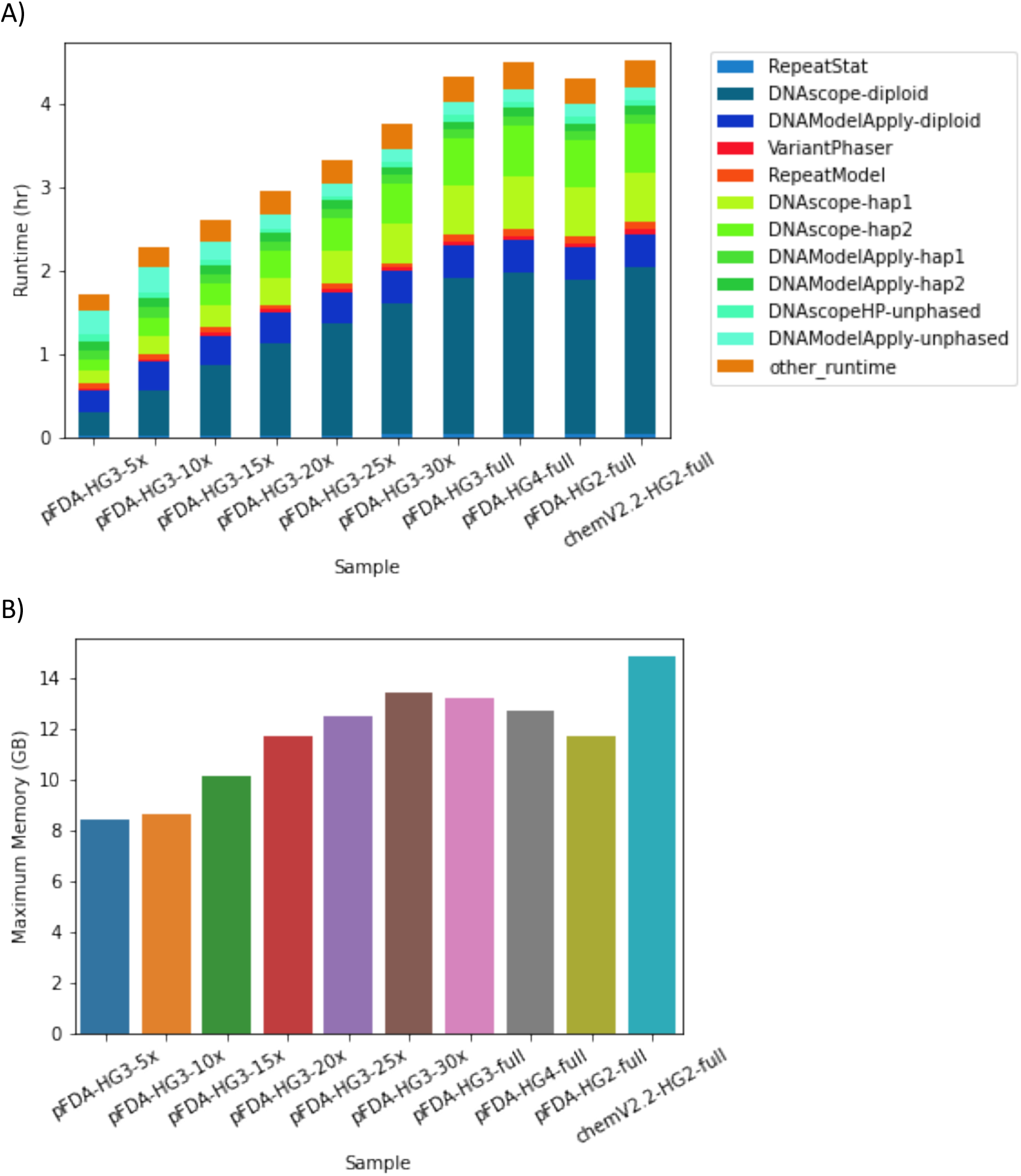
Runtime and maximum memory usage of the DNAscope LongRead pipeline. (A) Runtime of the DNAscope LongRead pipeline by pipeline stage on a 32-core Intel^®^ Xeon^®^ server. Stages are ordered chronologically in the pipeline. Stages in the first pass of variant calling are shades of dark blue, variant phasing stages are red, and stages in the second pass of variant calling are shades of green. The “other runtime” stage is short VCF manipulation commands. (B) Maximum memory usage of the pipeline by sample. Higher coverage samples generally have a higher memory usage during variant calling.

With high-coverage samples, the DNAscope LongRead pipeline spends approximately 54% of its overall runtime in the first stage of the pipeline, including diploid repeat model calibration, diploid variant candidate generation, and diploid SNV calling. Variant phasing and phased repeat model calibration is very efficient, consuming only about 3% of the overall runtime. Later passes of haploid and diploid variant calling consume approximately 36% of the overall runtime. The remaining 7% of runtime is consumed by VCF manipulation operations (Figure 4A).

The DNAscope LongRead pipelines in this study used a maximum memory of 14.8 GB for the X coverage sample (Figure 4B). Minimum memory usage was 8.39 GB for the 5x sample and overall memory usage was correlated with sample coverage. Within the pipeline, variant candidate generation (DNAscope) and variant phasing (VariantPhaser) commands had the highest memory usage.

The DNAscope LongRead pipeline provides scalable and highly efficient germline variant calling from PacBio HiFi reads. Variant calling from the 30x sample took 120.3 core-hours on our machine, and overall runtime increased proportionally with increasing coverage. Approximate On-Demand compute cost of germline variant calling for a representative 30x sample would be less than $6 in the AWS US East (Ohio) region as of 12/13/2021.

## Discussion

In this paper we introduce DNAscope LongRead, a highly accurate, efficient, robust and scalable pipeline for germline variant calling from PacBio HiFi reads. We find that the DNAscope LongRead pipeline provides a 15% reduction in mean errors from the Precision FDA Truth Challenge V2 winning variant calling pipeline for all benchmark regions, and for most samples with 30x (or greater) coverage, typically calls fewer than 10,000 total errors when evaluated against the Genome in a Bottle v4.2.1 benchmark dataset. Through serial downsampling of a 35x sample, we find SNV and indel variant calling accuracy extends well to lower coverages with a F1-score decreases of 0.00074 and 0.013, respectively, when moving from 35x to 20x. Variant calling is robust to changes in upstream data processing and retains high accuracy when evaluated across challenging genomic regions using novel benchmark datasets. The entire pipeline is efficiently implemented, performing analysis from inputs to the final variant dataset with a 30x sample in 120.3 core-hours with a peak memory usage of 13.44 GB. DNAscope LongRead is implemented using Sentieon 202112 a production ready pipeline for germline variant calling from PacBio HiFi reads.

## Methods

### Overview

The analysis described in this manuscript, including preprocessing, variant calling, truthset evaluation, and initial figure generation, were implemented as a Snakefile and run with snakemake version 6.3.0^18^. The complete snakemake pipeline is available at, https://github.com/DonFreed/dnascope-longread-manuscript/blob/main/scripts/pipeline.smk.

### Data preprocessing

HiFi reads were aligned to hs38 reference genome and sorted by genomic coordinate with pbmm2 version 1.4.0, which is a wrapper for minimap2 version 2.15. pbmm2 alignment was performed with the recommended settings, `--sort -j $threads --preset HiFi -c 0 -y 70`. The resulting BAM files were merged and converted to CRAM format using the Sentieon ReadWriter algo (version 202010.04).

Reads from the precisionFDA Truth Challenge V2 HG003 PacBio HiFi sample were then serially downsampled with the subsampling functionality in ’samtools view’ version 1.13. Expected coverage for each sample was confirmed with the Sentieon CoverageMetrics algo, version 202010.04.

### Variant calling using the Sentieon LongRead pipeline

Variants were called from the aligned (and optionally subsampled) reads using the Sentieon LongReads pipeline (version Beta0.4). Prior to variant calling, the input CRAM file was copied onto the machine’s local SSD. The DNAscope LongRead pipeline was run with a dbSNP version 146 VCF file, and an MHC BED file containing the hg38 coordinates of the MHC region (chr6:28510020-33480577). A BED file was used with the ’-b’ argument to restrict variant calling to diploid chromosomes in the sample (autosomes for males and autosomes and the X chromosome for females). The DNAscope LongRead pipeline calls `sentieon`, `bcftools`, and `bedtools` executables from the user’s PATH and was used with `sentieon` version 202010.04, `bcftools` version 1.11, and `bedtools` v2.25.0.

Variant calling was performed on a dedicated 32-virtual core Intel^®^ Xeon^®^ server with 64GB of memory and two 2TB dual stripped SSDs combined with RAID0 for local file storage.

### Variant callset evaluation

Variants called by the DNAscope LongRead pipeline were evaluated against either version v4.2.1 of the GIAB truthset^19^ or v1.00 of the GIAB Challenging Medically-Relevant Genes benchmark^17^ using hap.py^20^ at commit c69f402 and hap.py was configured to use rtgtools vcfeval version 3.8.2 as a variant comparison engine^21^. Hap.py was run with the additional parameters, ’--no-decompose --no-leftshift’. Hap.py’s variant stratification was used with the GA4GH stratification regions v2.0 to perform a stratified analysis^22^.

## Code Availability

Scripts for reproducing the described analysis can be found at https://github.com/DonFreed/dnascope-longread-manuscript

## Data Availability

The hs38 reference genome used in this work is available for download from the NCBI’s ftp site: ftp://ftp.ncbi.nlm.nih.gov/genomes/all/GCA/000/001/405/GCA_000001405.15_GRCh38/seqs_for_alignment_pipelines.ucsc_ids/GCA_000001405.15_GRCh38_full_analysis_set.fna.gz. The DNAscope LongRead pipeline used an optional dbSNP VCF file, which is available from the Broad’s FTP resource bundle at, ftp://ftp.broadinstitute.org/bundle//hg38/dbsnp_146.hg38.vcf.gz. GIAB v4.2.1 and CMRG v1.00 truthsets (VCF and BED files) are available for download from the GIAB consortium’s FTP site, ftp://ftp-trace.ncbi.nlm.nih.gov/giab/ftp/release/. Stratification regions used in the hap.py stratified analysis can be found at the GIAB consortium’s FTP site, ftp://ftp-trace.ncbi.nlm.nih.gov/giab/ftp//release/genome-stratifications/v2.0/GRCh38.

The PacBio HiFi data sequenced as part of the PrecisionFDA Truth Challenge V2 can be downloaded from the PrecisionFDA Truth Challenge V2 webpage, https://precision.fda.gov/challenges/10. The Chemistry v2.2 dataset is available in NCBI BioProject PRJNA832505.

## Acknowledgments

We would like to thank R. Aldana and B. Gallagher for helpful discussions regarding the proposed experiments. We would like to thank the investigators of the Genome in a Bottle consortium for their efforts in developing community resources for benchmarking. We would also like to thank the organizers of the precisionFDA Truth challenges.

## Supplementary Figures

**Supplementary Figure 1.**
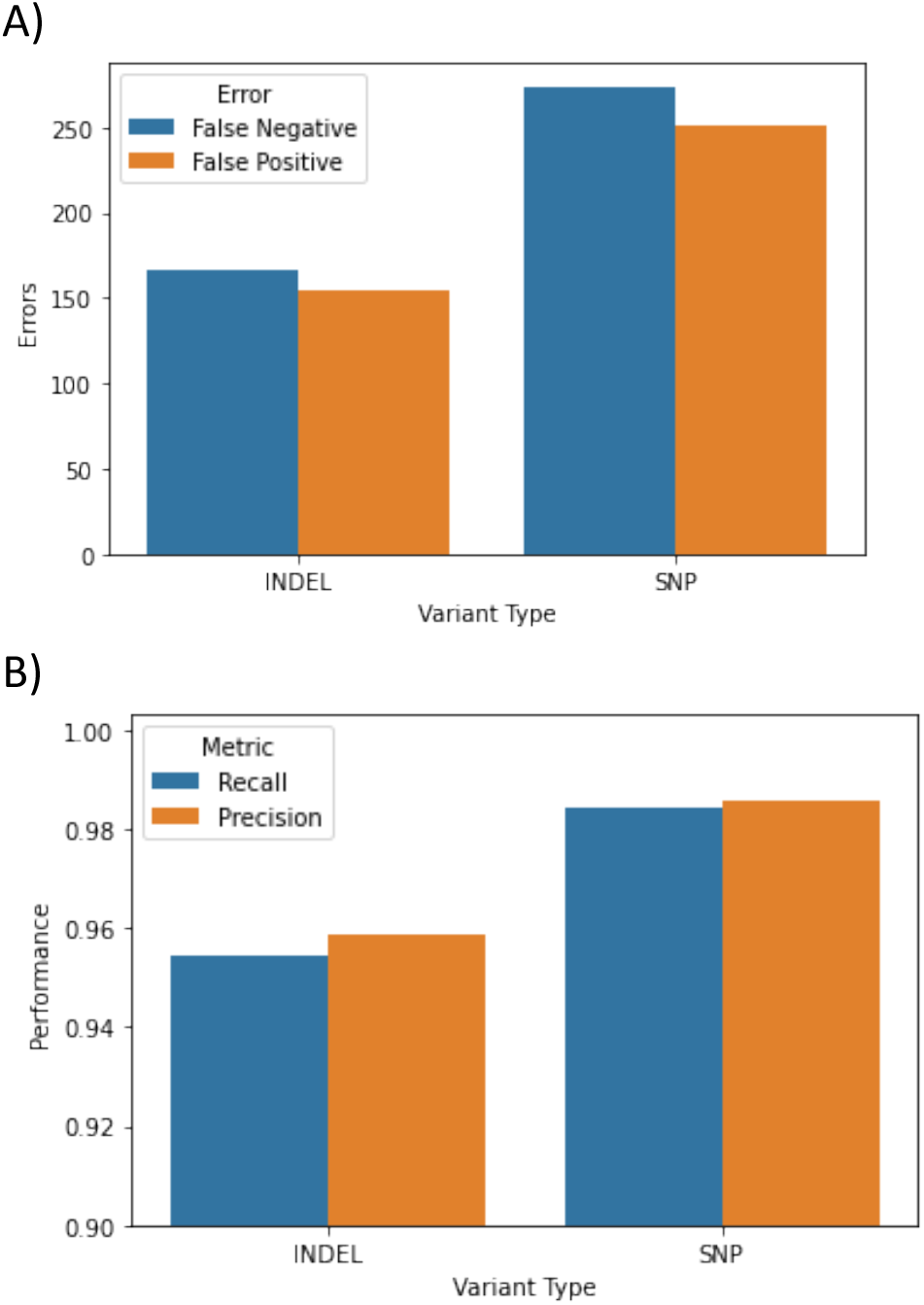
Accuracy of the PrecisionFDA Truth V2 sample evaluated on the Challenging Medically Relevant Genes benchmark. Accuracy of the DNAscope LongRead pipeline when evaluated with the Challenging Medically Relevant Genes benchmark (CMRG). The CRMG benchmark extends to more difficult regions outside of the v4.2.1 GIAB benchmark regions using assembly-based approaches. (A) Number of errors. (B) Precision and recall.

**Supplementary Figure 2.**
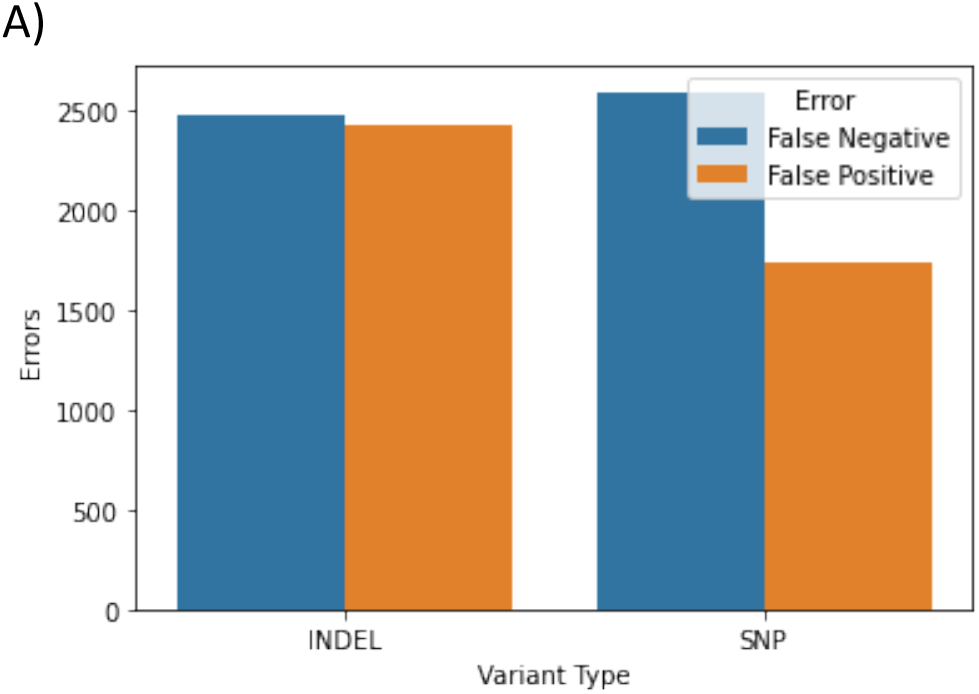
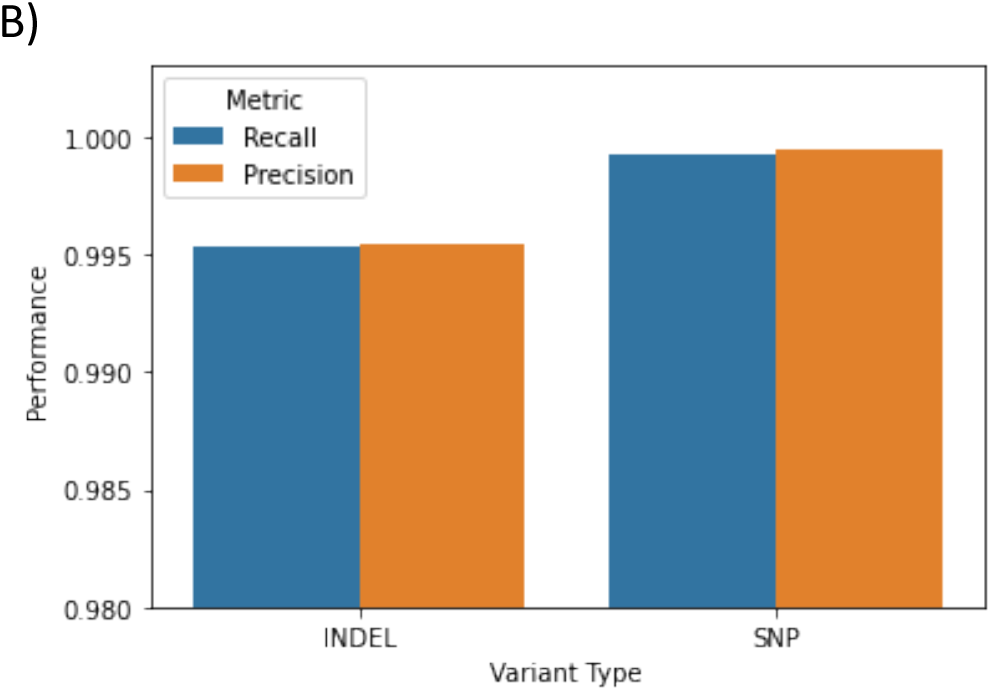
Accuracy of the Chemistry v2.2 sample evaluated on the GIAB v4.2.1 truthset. Accuracy of the DNAscope LongRead pipeline with the chemistry v2.2. sample when evaluated with the GIAB v4.2.1 benchmark. The DNAscope LongRead pipeline maintains high accuracy with the novel sequencing chemistry. (A) Number of errors. (B) Precision and recall.

## Supplementary Tables

**Supplementary Table 1. Evaluation of the highest scoring precisionFDA submissions and DNAscope LongRead on the precisionFDA Truth Challenge V2 PacBio HiFi samples**.

**Supplementary Table 2. Evaluation of the DNAscope LongRead pipeline with serial downsampling of the HG003 precisionFDA Truth Challenge V2 PacBio HiFi sample**.

**Supplementary Table 3. Samples used during training of the DNAscope LongRead machine learning model**.

**Supplementary Table 4. DNAscope LongRead accuracy on the PacBio Chemistry v2.2 sample**.

**Supplementary Table 5. Runtime benchmarks for the samples processed in this study**. Runtimes are in hours.

**Supplementary Table 6. Memory benchmarks for the samples processed in this study**. Maximum memory usage (max_vms) in gigabytes.

